# A language-familiarity effect on the recognition of computer-transformed vocal emotional cues

**DOI:** 10.1101/521641

**Authors:** Tomoya Nakai, Laura Rachman, Pablo Arias, Kazuo Okanoya, Jean-Julien Aucouturier

## Abstract

People are more accurate in voice identification and emotion recognition in their native language than in other languages, a phenomenon known as the language familiarity effect (LFE). Previous work on cross-cultural inferences of emotional prosody has left it difficult to determine whether these native-language advantages arise from a true enhancement of the auditory capacity to extract socially relevant cues in familiar speech signals or, more simply, from cultural differences in how these emotions are expressed. In order to rule out such production differences, this work employed algorithmic voice transformations to create pairs of stimuli in the French and Japanese language which differed by exactly the same amount of prosodic expression. Even though the cues were strictly identical in both languages, they were better recognized when participants processed them in their native language. This advantage persisted in three types of stimulus degradation (jabberwocky, shuffled and reversed sentences). These results provide univocal evidence that production differences are not the sole drivers of LFEs in cross-cultural emotion perception, and suggest that it is the listeners’ lack of familiarity with the individual speech sounds of the other language, and not e.g. with their syntax or semantics, which impairs their processing of higher-level emotional cues.

## Introduction

In everyday life, we interact socially with a variety of other people, some of them from cultural groups with which we have had limited previous encounters. A large body of psychological evidence shows that such familiarity, or lack thereof, with the cultures of others crucially affects how we process social signals such their facial or vocal features. People recognize faces of their own race more accurately than faces of other races (other-race effect; Shapiro & Penrod, 1986; Meissner & Brigham, 2001), and identify speakers of their native language better than speakers of other languages (language-familiarity effect or LFE; Perrachione et al., 2011; Fleming et al., 2014). Even within a given cultural or language group, familiarity with e.g. a given speaker’s voice facilitates how his spoken words are remembered (Pisoni, 1993), or how ambiguous prosodic cues are processed semantically (Chen et al., 2016). These effects are thought to result primarily from perceptual learning: differential exposure wraps an observer’s perceptual space to better accommodate distinctions that are informative to discriminate common, in-group items, with the result that comparatively less-common, out-group items are encoded in a less efficient manner (Valentine, 1991; Furl et al., 2002). Such perceptual warping is manifest e.g. in the fact that listeners rate pairs of speakers of their own language as more dissimilar than pairs of speakers of the other language (Fleming et al., 2014), or that young infants can discriminate sounds from all languages equally well before six months of age, but develop a native-language advantage by about 6-12mo (Kuhl et al., 1992; Johnson et al., 2011).

One cognitive domain in which language-familiarity effects may be particularly prevalent is that of cross-cultural inferences of emotional prosody (Scherer et al., 2001; Thompson & Balkwill, 2006; Pell et al., 2009; Sauter et al., 2010). Judging whether a particular speech utterance is unusually high or low, a given phoneme bright or dark, whether a specific pitch inflection is expressive or phonological (Scherer & Oshinsky, 1977; Juslin & Laukka, 2003) all would appear to be advantaged, or conversely impaired, by acquired auditory representations optimized for the sounds of one language or another. Most cross-cultural data indeed reveal an in-group advantage for identifying emotions displayed by members of the same rather than a different culture (Scherer et al., 2001; Thompson & Balkwill, 2006; Pell et al., 2009; see Elfenbein & Ambady, 2002 for a review), showing that vocal emotion recognition is governed by both language-independent (universal) and language-specific processes. However, it has been difficult to conclusively determine to what extent such differences arise from language-familiarity effects in the sense of the above, or more generally from learned cultural differences in how these emotions are expressed (Elfenbein & Ambady, 2002). For instance, LFEs would predict better cross-cultural recognition with increasing language similarity, but such evidence is mixed: Scherer and colleagues (Scherer et al., 2001) found Dutch listeners better at decoding German utterances than listeners of other, less similar European and Asian languages, but other studies found no differences in e.g. how Spanish listeners identified vocal emotions from the related English and German languages, and the unrelated Arabic (Pell et al., 2009), or how English listeners processed utterances in the related German language and the unrelated Tagalog of the Philippines (Thompson & Balkwill, 2006). Most critically, because most of such cross-cultural investigations use stimuli recorded by actors of each culture, differences in the agreement levels across groups may simply arise because of cultural differences in emotional expressions. If the last author, a Frenchman, has difficulties processing emotional cues spoken by the first author, is it because his auditory representations of the Japanese phonetic inventory are poorly differentiated (see e.g. Dupoux et al., 1999), or because one simply does not use the same cues to express *joy* in Japanese and in French (see e.g. Kitayama et al., 2006)? To progress on this debate, it would be necessary to employ emotional voice stimuli which, while being recognized as culturally-appropriate emotional expressions in different languages, utilize *exactly* the same prosodic cues in *exactly* the same manner (e.g. a 50-cent pitch increase on the second syllable), something which is at best impractical with vocal actors.

To this aim, we used a novel software platform (DAVID, Rachman et al., 2017) to apply a fixed set of programmable emotional transformations to prerecorded voice samples in two languages (French and Japanese), and performed cross-cultural emotion recognition experiments to assess how identical emotional cues were processed in both languages by speakers of both cultures. In brief, DAVID alters the acoustic features of original voices by combining audio effects of pitch shift, vibrato, inflection and filtering, to the result of modifying the emotional content of the original voice (Fig. 1). In a first study with DAVID, authors applied real-time emotion transformations on participants’ voices while they were talking. Transformed voices were perceived as natural emotional expressions, to the point that most participants remained unaware of the manipulation, but they nevertheless affected participants’ emotional states and skin conductance responses (Aucouturier et al., 2016). In a follow-up study, authors tested the effect of DAVID in the multiple languages, finding that transformed emotions were recognized at above-chance levels when applied to either French, English, Swedish, or Japanese utterances, and with a naturalness comparable to authentic speech (Rachman et al., 2017). These studies provide basic evidence that DAVID emotion transformations can be applied to different languages.

**Figure 1:**
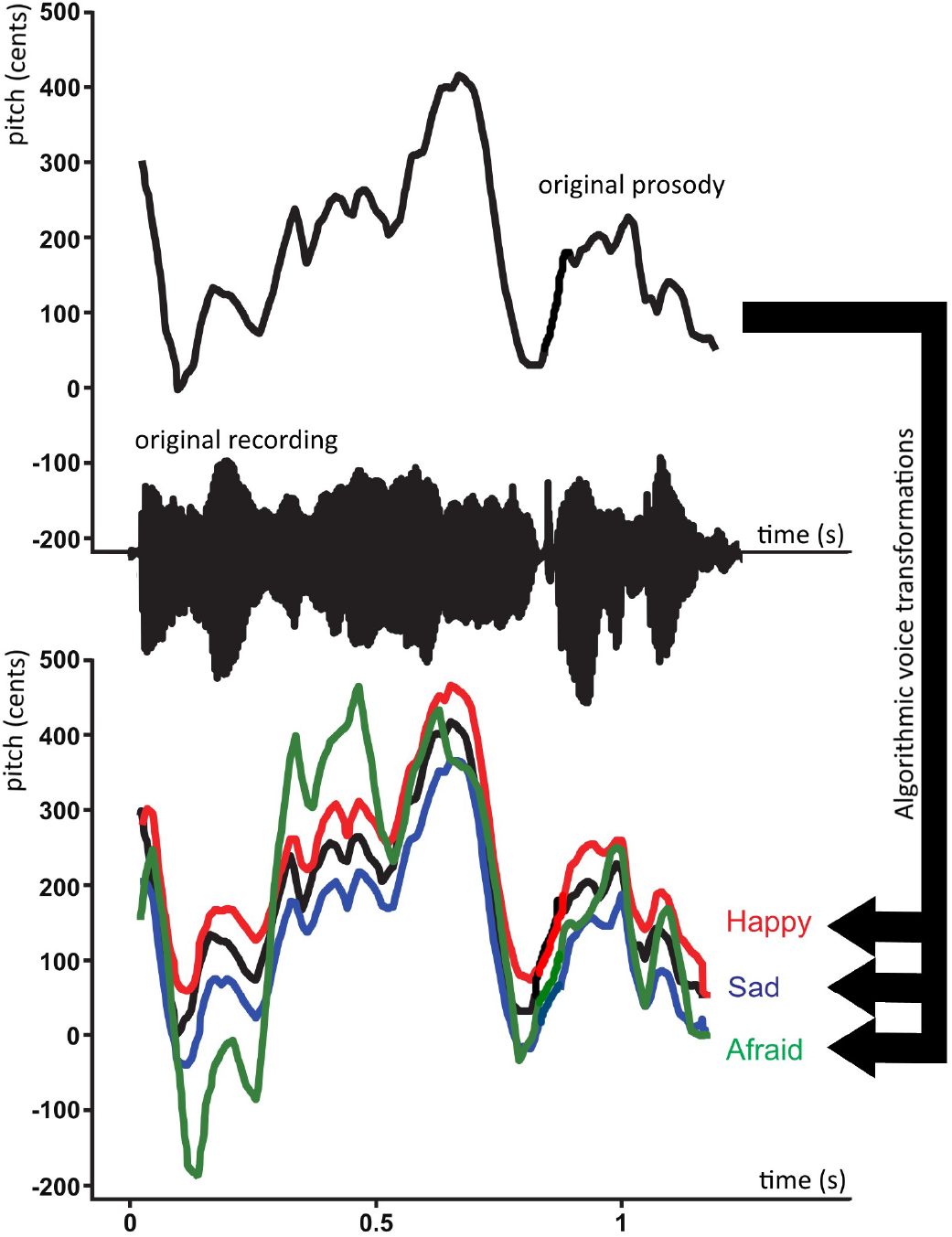
Illustration of the algorithmic voice transformations used in the study. A single recording of a French female speaker saying *"J’ai oublié mon pardessus"* (I forgot my jacket) is manipulated with the DAVID voice transformation plateform to make it sound more happy, sad or afraid. Solid black: time series of pitch values in the original recording estimated with the SWIPE algorithm Camacho & Harris (2008). Red, blue, and green lines: pitch of manipulated audio output in the Happy, Sad, and Afraid transformations, respectively. The speech waveform of the unmodified recording is shown on the x-axis. Pitch values on y-axis are normalized to cents with respect to mean frequency 200 Hz.

In the present study, we manipulated both Japanese (JP) and French (FR) voices with the same set of parametric transformations by DAVID, so as to display emotions of happiness, sadness and fear/anxiety. This procedure excludes the possible effect of different emotion expressions and individual or cultural variability of recordings, because emotion categories of two languages are produced exactly in the same manner (i.e., with the same algorithmic parameters). Twenty-two native JP and twenty-two native FR speakers listened to pairs of such computer-manipulated voices (one neutral, one manipulated), in both languages, and force-choice categorized the emotion of the second voice. To test which linguistic component contributed to such LFE, we further prepared syntactically/semantically modified sentences (see Methods for the detailed explanation), as well as time-reversed voices and sentences in a non-familiarized language for both participant groups (Swedish sentences).

## Methods

### Participants

Twenty-two native Japanese (JP) speakers (9 female, M=19.7) and 22 French (FR) speakers (12 female, M=23.6) participated in the current experiment. None of the JP speakers had ever learned French, and none of the FR speakers had ever learned Japanese. Experiments with the Japanese speakers were conducted in the University of Tokyo (Japan), while those with the French speakers were conducted at the INSEAD/Sorbonne-Université Behavioural platform in Paris (France). Volunteers were recruited through local databases and mailing lists in the respective countries and were financially compensated for their participation. Two French participants were excluded because they did not satisfy language requirements (not native FR speakers). Furthermore, one Japanese speaker was excluded because they claimed that they could hear the auditory stimuli only from one side of the headphone.

### Ethics

All experiments were approved by the Institut Européen d’Administration des Affaires (INSEAD) IRB, as well as by the departmental review boards of the University of Tokyo, Komaba. All participants gave their informed consent and were debriefed about the purpose of the research immediately after the experiment.

### Stimuli

We prepared four normal JP sentences and four normal FR sentences, translated from the same four semantically-neutral English sentences (see Table 1 and Russ et al., 2008). Jabberwocky variants (Hahne & Jescheniak, 2001) of the eight sentences were created by replacing each content word in the normal sentences to a pseudo-word with the same number of syllables, without changing functional words (e.g. JP: *Uwagi wo wasureta* (I forgot my jacket) −> *Etaki* wo morushita\**). Jabberwocky sentences did not have any meaning (or content), but still maintained the grammatical structures of their original languages. Shuffled Jabberwocky sentences were produced by changing the word order of the corresponding Jabberwocky sentences so that they did not maintain any grammatical structures of their original languages (e.g. JP:Etaki *wo morushita* −> *Wo* morushita* etaki\**; see Table 1). The number of syllables was balanced across the two languages (7.25 ± 1.5 [mean ± SD] for the normal JP stimuli, and 7.5 ± 1.3 for the normal FR sentences), and the Jabberwocky and shuffled variants of each sentence had the same number of syllables as the original version, in both languages.

**Table 1:** Original sentences

| JP sentences | FR sentences | English translations |
| --- | --- | --- |
| **Normal** |  |  |
| Uwagi wo wasureta | J'ai oublié mon pardessus | I forgot my jacket |
| Kaigi ni itta | Je suis allé à la réunion | I attended the meeting |
| Hikouki wa ippai | L'avion est complet | The airplane is full |
| Hyoumen wa nameraka | La surface est lisse | The surface is slick |
| **Jabberwocky** |  |  |
| Etaki wo morushita | J'ai odrié mon tarpodu |  |
| Goito ni etta | Je suis ijé à la boussion |  |
| Komeubi wa ottai | L'ation est bondret |  |
| Bouren wa soniyaka | La borpaque est nitte |  |
| **Shuffled** |  |  |
| Wo morushita etaki | Odrié j'ai tarpodu mon |  |
| Etta ni goite | Boussion ijé la je à suis |  |
| Komeubi ottai wa | L'est ation bondret |  |
| Bouren soniyaka wa | La est nitte borpaque |  |

French and Japanese normal, jabberwocky, and shuffled sentences used in this study. English translations are added for clarification, but were not included in this study.

In each language, one male and one female speaker recorded the four normal, four jabberwocky, and four shuffled sentences corresponding to their native language. We then used the recordings of the normal JP and FR sentences to create 16 additional reversed recordings (four sentences, two speakers, in each language), by playing them backwards. The recordings took place in a sound-attenuated booth, using Garage-Band software (Apple Inc.) with a headset microphone. In addition, we used four male and four female normal Swedish (SE) recordings from a previous study (Rachman et al., 2017). All stimuli were normalized in length (1.5 s) using a phase vocoder algorithm (AudioSculpt, IRCAM) and in root-mean-square intensity.

Finally, all the above recordings (incl. reverse JP, reverse FR, and normal SE) were processed with DAVID and transformed into happy, sad, and afraid variants, resulting in 96 manipulated recordings for both JP and FR (3 emotions × 4 sentences × 4 conditions × 2 speakers), and 24 manipulated SE recordings. (In particular, note that reversed recordings were manipulated after, and not before, being reversed.)

### Audio manipulation algorithm

Recordings were manipulated with the software platform DAVID (Rachman et al., 2017), using predetermined pitch shift, vibrato, inflection, and filtering transformations designed to evoke happy, sad, and afraid expressions.

Pitch shifting denotes the multiplication of the pitch of the original voice signal by a constant factor α (e.g. + 50 cents, a 2.93% change of F0). Vibrato is a periodic modulation of the pitch of the voice, occurring with a given rate and depth. Vibrato is implemented as a sinusoidal modulation of the pitch shift effect, with a rate parameter (e.g. 8 Hz), a depth (e.g. + 40 cents) and a random variation of the rate (e.g. 30% of the rate frequency). Inflection is a rapid modification of the pitch at the start of each utterance, which overshoots its target by several semitones but quickly decays to the normal value. DAVID analyzes the incoming audio to extract its root-mean-square (RMS), using a sliding window. When the RMS reaches a minimum threshold, the system registers an attack, and starts modulating the pitch of each successive frame with a given inflection profile. Finally, filtering denotes the process of emphasizing or attenuating the energy contributions of certain areas of the frequency spectrum (e.g., a high-shelf filter with a cut-off frequency at 8000 Hz, +9.5 dB per octave). Full details of these algorithms can be found in Rachman et al. (2017).

Table 2 describes the transformation parameter values used in this study to simulate the emotions of happy, sad, and afraid, which were applied identically to both Japanese and French-language stimuli. These values were based on previous studies (Aucouturier et al., 2016; Rachman et al., 2017) that validated the recognizability and naturalness of the transformations in both JP and FR. Examples of the manipulations are illustrated in Figure 1, and in supplementary audio files.

**Table 2:**
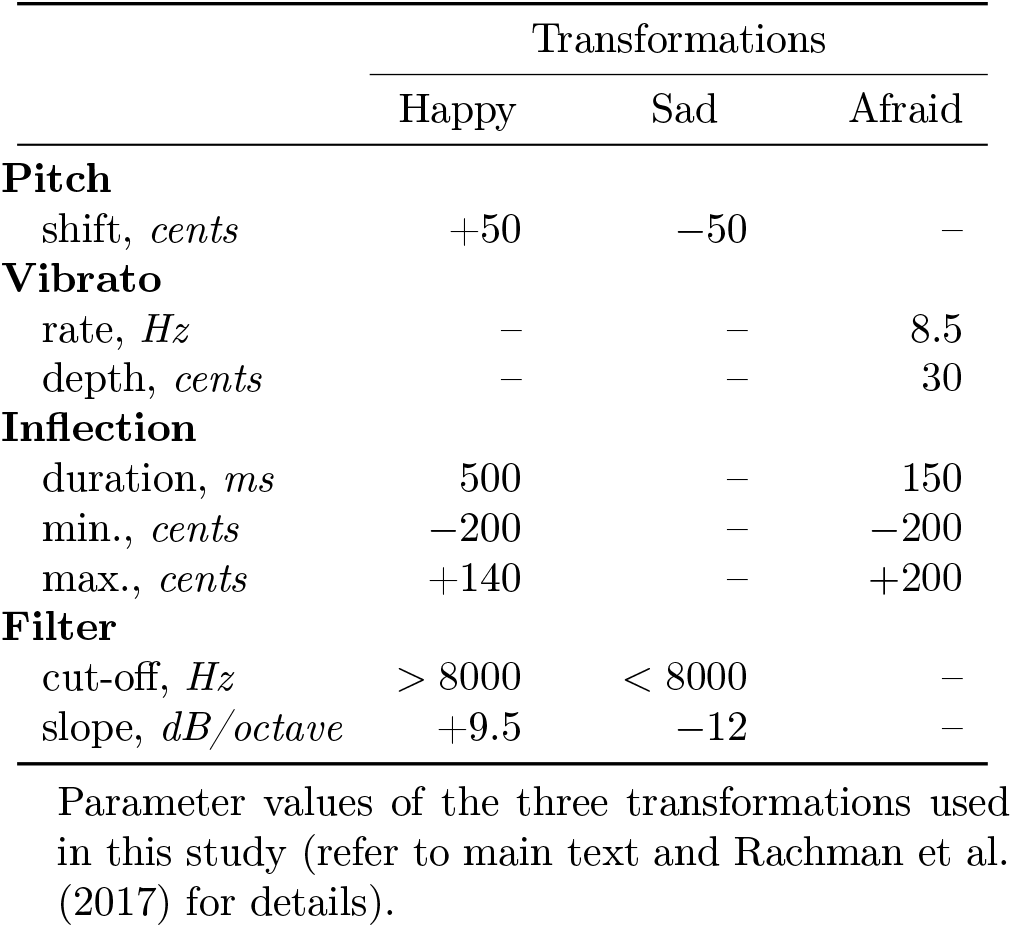
Deviant parameter values

|  | Transformations |  |  |
| --- | --- | --- | --- |
|  | Happy | Sad | Afraid |
| **Pitch** |  |  |  |
| shift, *cents* | +50 | -50 | - |
| **Vibrato** |  |  |  |
| rate, *Hz* | - | - | 8.5 |
| depth, *cents* | - | - | 30 |
| **Inflection** |  |  |  |
| duration, *ms* | 500 | - | 150 |
| min., *cents* | -200 | - | -200 |
| max., *cents* | +140 | - | +200 |
| **Filter** |  |  |  |
| cut-off, *Hz* | > 8000 | < 8000 | - |
| slope, *dB/octave* | +9.5 | -12 | - |

Parameter values of the three transformations used in this study (refer to main text and Rachman et al. (2017) for details).

### Procedure

In each trial, participants listened to pairs of utterances of the same sentence by the same speaker. The first utterance was always the neutral recording and the second utterance was either the same recording unprocessed (neutral condition) or processed with one of the emotional transformations (happy, sad, afraid). After hearing the two utterances (ISI=0.7-1.3 s), participants were instructed to answer whether the second utterance sounded happy, sad, afraid, or neutral by pressing one of four keys (“S”, “D”, “F”, and “G”) with their left fourth, third, second, and first finger, respectively. The next trial started when participants pressed the “ENTER” key with their right first finger. All participants were presented with the 96 JP pairs (4 sentences × 2 speakers × 4 conditions × 4 emotions), 96 FR pairs, and 24 SE pairs, randomized across participants in one single block. The correspondence of keys and response categories was randomized across participants. Visual stimuli were displayed on a laptop-computer screen, and the voice stimuli were presented through closed headphones. Stimulus presentation and data collection were controlled using PsychoPy toolbox (Peirce, 2007). Response categories in the recognition task for the French group used the English terms (happy, sad, afraid) instead of the French equivalents, but were defined in the instructions using the equivalent French terms. Response categories used in Japanese group were presented in Japanese terms.

### Data analysis

We computed mean hit rate of the three emotion categories (happy, sad, and afraid) for each of the nine conditions (JP, FR: normal, jabberwocky, shuffled, reversed; SE: normal). To take response bias into account, we also calculated unbiased hit rates for each participant (Wagner, 1993).

## Results

Both participant groups recognized the three emotional transformations above chance level when applied to normal speech of their native language (Fig.2.A). For FR participants, average hit rate over the three emotion categories was significantly larger than chance level (25 %) in normal FR sentences (*p* = .0024, *α_Bonfer_*. = .0083). For JP participants, hit rate was larger than chance for both normal JP and normal FR (*p* < .001). Hit rates for the normal SE were around chance level for both participants groups. In both participant groups, hit rates on native language were at comparable levels (FR:.37; JP:.44; *t*(39) = 1.40,*p* = .85, *d* = 0.44). These results confirmed both that participants had good emotion recognition abilities in their native language, and that the emotional transformations used in this study are appropriate (and of comparable discriminability) in both languages.

**Figure 2:**
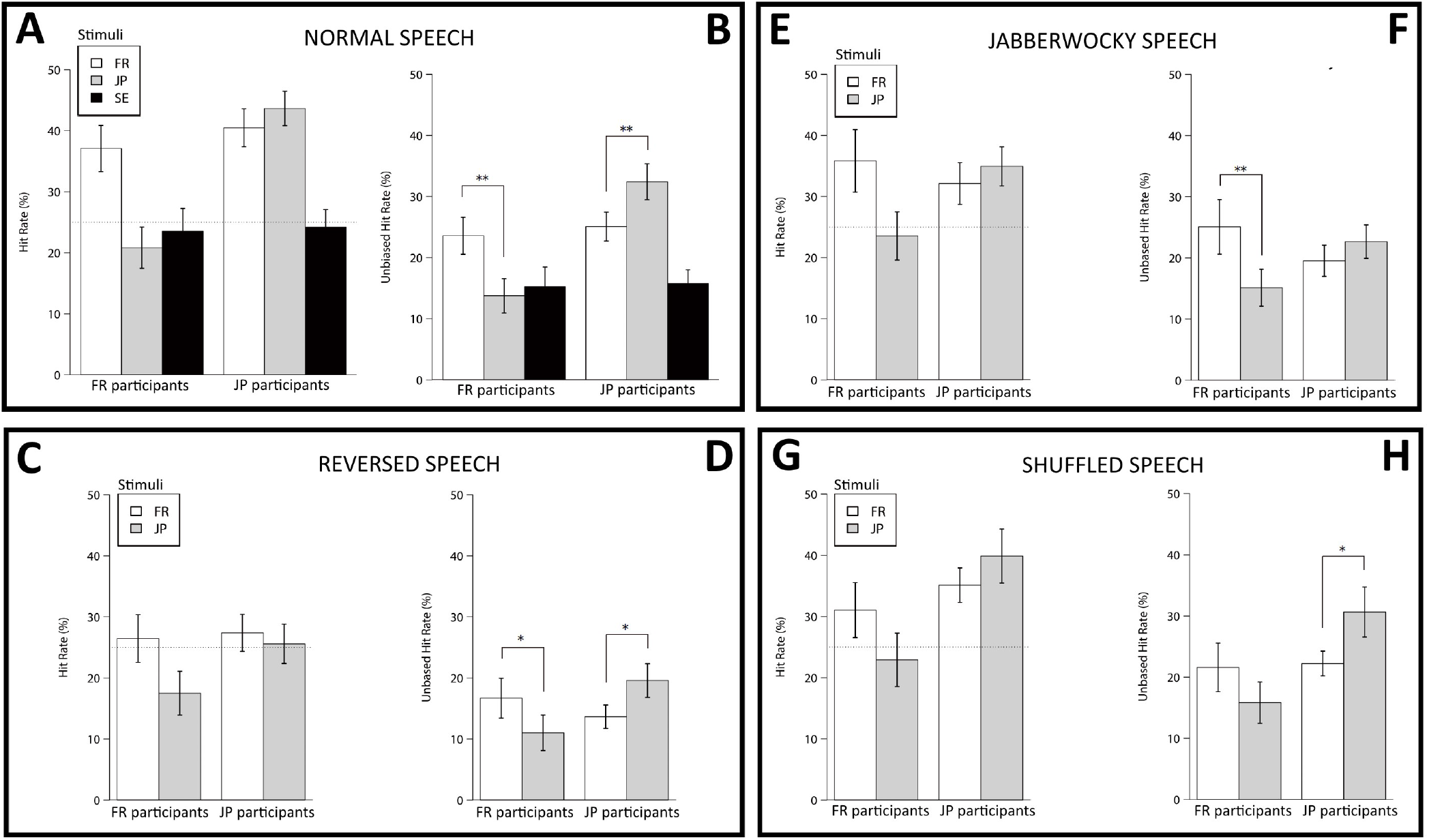
Biased (A,C,E,G) and unbiased (B,D,F,H) hit rates averaged over the 3 non-neutral emotion categories, grouped by normal, reversed, jabberwocky and shuffled speech conditions. In the normal condition, in addition to the results of French-language (in white) and Japanese-language (in gray) sentences, those of Swedish-language sentences (in black) are also shown. \**p* < .05. \*\**p* < .01. Error bar, SEM.

To test for language-familiarity effects, we then examined unbiased hit rates. A two-way repeated measures analysis of variance (rANOVA) of Language × Participant group showed a significant interaction (*F*(1,39) = 22.02, *p* < .001, 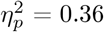), as well as a significant main effect of participant group (*F*(1,39) = 8.33, *p* = .006, 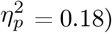). Post-hoc *t*-tests (*α_Bonfer_*. = 0.025) showed that FR participants were able to detect the emotional categories in normal FR sentences significantly better than in normal JP sentences (*t*(1,19) = 3.60, *p* < .001, *d* = 0.75), and JP participants could detect emotions in normal JP sentences significantly better than in normal FR sentences (*t*(1,20) = 3.00, *p* = .003, *d* = 0.60; see Fig. 2.B). These results indicate a clear LFE.

This LFE was observed in a quasi-identical manner in all three degraded stimulus conditions. When participants were tasked to decode the same emotional cues on reversed JP and FR stimuli, average hit rate was around chance level for both participant groups (*α_Bonfer_*. = 0.0125, Fig. 2.C). However, unbiased hit rates showed a significant interaction between Language and Participant group (*F*(1,39) = 14.05, *p* < .001, 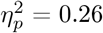), with FR participants better in reverse FR than reverse JP sentences (*t*(1,19) = 2.53, *p* = .010, *d* = 0.41), and JP participants better in the reverse JP than reverse FR sentences (*t*(1,20) = 2.78, *p* = .006, *d* = 0.54; Fig. 2.D).

Average hit rate of the FR participants was at chance level for both jabberwocky and shuffled sentences *α_Bonfer_*. = 0.0125, while JP participants showed better-than-chance performance in jabberwocky JP, shuffled JP, and shuffled FR sentences (*ps* = .003, .0009, .002, respectively; Fig. 2.E-G). Unbiased hit rates again showed a significant interaction of Language × Participant group in both jabberwocky sentences (*F*(1,39) = 12.06, *p* = .001, 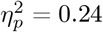), and shuffled sentences (*F*(1,39) = 9.37, *p* = .004, 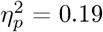). FR participants were better in the jabberwocky FR than jabberwocky JP sentences (*t*(1,19) = 3.23, *p* = .002, *d* = 0.58; Fig. 2.F), and JP participants were better in shuffled JP than shuffled FR sentences (*t*(1,20) = 2.32, *p* = .016, *d* = 0.57; Fig. 2.H). Language difference was marginally significant in jabberwocky sentences for JP participants (*t*(1,19) = 2.04, *p* = .028, *d* = 0.35), but not in shuffled sentences for FR participants (*t*(1,20) = 1.42, *p* = .086, *d* = 0.26).

Finally, we analyzed the three emotion categories (happy, sad, afraid) separately by averaging all of the four sentence types (normal, reverse, jabberwocky, shuffled), as all of these conditions showed an interaction effect of language and participant group (Fig. 3). For FR participants, the LFE affected all three emotions identically. A 2-way rANOVA of Emotion category × Language type showed a significant main effect of Emotion category (*F*(1,19) = 22.97, *p* < .001, 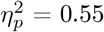), as well as a significant main effect of Language type (*F*(2,38) = 3.63, *p* = .003, 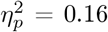), but no interaction between emotion and language. For JP participants however, we found a significant main effect of Emotion type (*F*(1,20) = 28.05, *p* = .003, 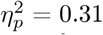), a main effect of Language type(*F*(2,40) = 8.78, *p* < .001, 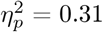), and an interaction (*F*(2,40) = 9.45, *p* < .001, 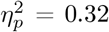). Further analyses showed simple main effects of Language type in Happy and Afraid emotions (both *p* <.001), but not in Sad emotion (*p* = .92).

**Figure 3:**
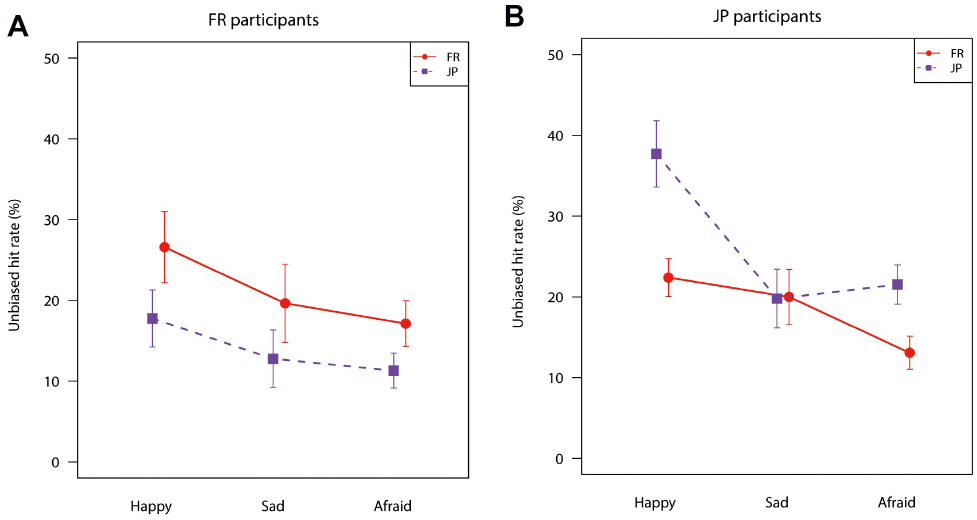
Unbiased hit rates for each emotion category for both FR (A) and JP participants (B), averaged across normal, reversed, jabberwocky, and shuffled conditions. Red solid line shows unbiased hit rat of FR sentences, while blue dashed line shows that of JP sentences. Error bar, SEM.

## Discussion

Previous work on cross-cultural emotion recognition had left it difficult to determine whether native-language advantages arise from language-familiarity effects of the sort seen with speaker or face recognition, or from cultural differences in how these emotions are expressed. In order to rule out such production differences, this work employed algorithmic voice transformations to create pairs of stimuli in the French and Japanese language which differed by exactly the same amount of prosodic expression. Even though the cues were strictly identical in both languages, they were better recognized when participants processed them in their native language. This advantage persisted in three types of stimulus degradation (jabberwocky, shuffled and reversed; Fig. 2).

These results provide univocal evidence that production differences (or “speaker dialects” Scherer et al., 2001) are not the sole drivers of in-group advantages in cross-cultural emotion perception. Even when we controlled e.g. happiness to be expressed with a precisely-calibrated pitch change of +50 cents in both languages, participants more accurately recognized such changes when they occurred in their native language. Critical to our manipulation is the fact that both groups reached identical performance on their normal native speech, showing that the computer-generated cues were equally discriminative in both languages. It could have been that computer manipulations differed in saliency depending on the phonetic characteristics of the language on which they were applied (e.g. vibrato needing relatively large consonant/vowel ratio, see Rachman et al., 2017), but this does not seem to be the case for the cues and the languages considered here. This native-language advantage can therefore only be explained by an interaction between the processes of encoding the linguistic or paralinguistic features of the familiar and non-familiar language and the processes responsible for extracting the emotional features important for the task - in short, by a language-familiarity effect of the kind already known for other-race face or speaker recognition (Meissner & Brigham, 2001; Perrachione et al., 2011). Emotional cue extraction may be facilitated in the case of native-language (e.g. better representation of what is phonological, and therefore better discrimination of what is incidental and expressive; see e.g. Dupoux et al., 1999), or negatively affected in the case of a non-familiar language (e.g. more effortful encoding, and therefore less resources available to process expressive cues; see e.g. Van Dillen & Derks, 2012), or both.

The percentage accuracy effect size of the native-language advantage in this study (FR: +7.2%, JP:+7.9% unbiased hit rate; Fig.3) was comparable with that of meta-studies of the in-group advantage in cross-cultural emotion recognition (+9.3%, Elfenbein & Ambady, 2002, p.216). The fact that studies included in this metaanalysis typically subsumed both speaker- and listener-level influences on emotion recognition, and in particular differences in how the emotions were displayed by actors cross-culturally, suggests that the perceptual LFE uncovered here is by no means a minor contributor in cross-cultural misperceptions, but may rather explain a large proportion of such effects.

When LFEs were suggested a possible driver of ingroup advantages in cross-cultural emotional voice perception (Pell & Skorup, 2008; Pell et al., 2009), it remained unclear at what level linguistic or paralinguistic processes interfered with emotional cue recognition: non-familiar stimuli differ both on lower-level auditory/phonological features, such as formant distributions and phoneme categories, mid-level prosodic representations, such as supra-segmental patterns for stress and prominence, but also higher-level syntax and semantics. Here, we contrasted normal stimuli with three degraded conditions, breaking semantics (grammatical sentences with non-words, or jabberwocky (Hahne & Jescheniak, 2001)), syntax (shuffled jabberwocky), and suprasegmental patterns (reversed speech, see e.g. Fleming et al., 2014), and found clear LFEs in all three conditions. This suggests that a low-level auditory/phonological basis for the effect, i.e. that it is the listeners’ lack of familiarity with the individual speech sounds of the other language that primarily impairs their processing of higher-level emotional cues. This finding is consistent with LFEs in speaker recognition, which are preserved with reversed-speech (Fleming et al., 2014) but absent in participants with impaired phonological representations Perrachione & Wong (2007), but less intuitive in the case of emotional prosody, the processing of which is in well-known interaction with syntax (Eckstein & Friederici, 2006) and semantics (Kotz & Paulmann, 2007).

For the (biased) hit rates, JP participants could recognize emotional cues in normal and shuffled FR sentences well above chance level, while the hit rates of FR participants for the JP were not. Although LFE is found for both participant groups, this asymmetry suggests that JP participants were more familiar to FR voices than FR participants to JP voices. One possible cause of such asymmetry is the effect of linguistic closeness of English and French as members of the Indo-European language family, and having influenced each other over history. While we confirmed that no JP participants had learned French before the experiment, Japanese students routinely learn English in the course of their education, and it is possible that familiarity with English facilitated emotion recognition in French (an effect also discussed in Scherer et al., 2001 in the case of German and Dutch).

Finally, we examined how LFEs differed by emotion category (Fig. 3). Two points deserve discussion. First, irrespective of listener language, happiness was more distinct in its recognizability across languages (M=+12.1% unbiased hit rate) than afraid (M=+7.1%) and sadness (M=+3.3%). This is consistent with previous meta-studies of in-group advantage in the voice modality (happiness: +17.5%; afraid: +12.9%; sadness: +7.6%; Elfenbein & Ambady, 2002, p.225), and confirms the general notion that expressions of happiness are a lot less universal in the vocal modality (Pell et al., 2009; Juslin & Laukka, 2003) than they are in the (overwhelmingly smile-driven) facial modality (Elfenbein & Ambady, 2002; Jack et al., 2012), possibly here because they rely on cues that require sufficiently accurate phonological representations of the target language to be extracted successfully. Interestingly, this would be the case, e.g., of the smiling oro-facial gesture which, universal as it may be in the visual domain, translates acoustically to fine formant-related variations of vowel spectra (Ponsot et al., 2017).

Second, we found that the performance of JP participants did not differ between native- and foreign-language stimuli for the sad emotion, although it did for happy and afraid. This may translate either a lack of precision of the sad emotion, or a lack of specificity of the sad response category, or both. It is possible that, while the computer-generated cues used here were generally appropriate for all emotions in both cultures, they were comparatively further away from the cultural norm of how sadness is expressed in the JP language. However, if these cues were wildly inappropriate for the JP culture, one would predict that JP participants would have similar difficulties recognizing such inappropriate cues when applied to FR stimuli, in which case we would see a main effect of language at a lower absolute level of recognition, but not a negation of the LFE for that emotion. Our data instead suggests that the processing of computer-generated cues was only impaired at a high-level of cultural expertise, indicating that any such ambiguity was masked when JP participants processed FR stimuli. This data is consistent with earlier reports of JP participants responding sad in response of happy JP stimuli more often than the other groups of participants (Rachman et al., 2017), and may result from different boundaries between emotional terms: it is possible that the cues manipulated in e.g. the happy effect spanned a larger proportion of the vocal expressions referred to as “sad” (*kanashimi*) in Japanese than the proportion of expressions referred to as “sad” (*triste*) in French. Future work could use reverse correlation paradigms such as those recently developed for the visual modality (Jack et al., 2012) to investigate how participants mentally represent typical prosody in these various emotional categories across cultures.

## Acknowledgments

T.N, L.R., K.O., and JJ.A. designed the experiment. L.R. validated the DAVID software, and P.A. recorded and processed the stimuli. T.N. collected the data. T.N. and JJ.A. analyzed and interpreted the data, using code contributed by L.R. T.N. and JJ.A. wrote the manuscript, with contributions from L.R., P.A. and K.O. This work was supported by ERC Grant StG 335536 CREAM and MEXT/JSPS KAKENHI Grant Number 4903, JP17H06380. All the data in this experiment was collected at the Centre Multidisciplinaire des Sciences Comportementales Sorbonne Université–Institut Européen d’Administration des Affaires (INSEAD). The authors thank Mael Garnotel who collected data in the French group. All data reported in the paper are available on request.

## References

Aucouturier, J.-J., Johansson, P., Hall, L., Segnini, R., Mercadié, L., & Watanabe, K. (2016). Covert digital manipulation of vocal emotion alter speakers’ emotional states in a congruent direction. Proceedings of the National Academy of Sciences, 113 (4), 948–953.

Camacho, A., & Harris, J. G. (2008). A sawtooth waveform inspired pitch estimator for speech and music. The Journal of the Acoustical Society of America, 124 (3), 1638–1652.

Chen, B., Kitaoka, N., & Takeda, K. (2016). Impact of acoustic similarity on efficiency of verbal information transmission via subtle prosodic cues. EURASIP Journal on Audio, Speech, and Music Processing, 2016 (1), 19.

Dupoux, E., Kakehi, K., Hirose, Y., Pallier, C., & Mehler, J. (1999). Epenthetic vowels in japanese: A perceptual illusion? Journal of experimental psychology: human perception and performance, 25 (6), 1568.

Eckstein, K., & Friederici, A. D. (2006). It’s early: event-related potential evidence for initial interaction of syntax and prosody in speech comprehension. Journal of Cognitive Neuroscience, 18 (10), 1696–1711.

Elfenbein, H. A., & Ambady, N. (2002). On the universality and cultural specificity of emotion recognition: a meta-analysis. Psychological bulletin, 128 (2), 203.

Fleming, D., Giordano, B. L., Caldara, R., & Belin, P. (2014). A language-familiarity effect for speaker discrimination without comprehension. Proceedings of the National Academy of Sciences, 111 (38), 13795–13798.

Furl, N., Phillips, P. J., & O’Toole, A. J. (2002). Face recognition algorithms and the other-race effect: computational mechanisms for a developmental contact hypothesis. Cognitive Science, 26 (6), 797–815.

Hahne, A., & Jescheniak, J. D. (2001). What’s left if the jabberwock gets the semantics? an erp investigation into semantic and syntactic processes during auditory sentence comprehension. Cognitive Brain Research, 11 (2), 199–212.

Jack, R. E., Garrod, O. G., Yu, H., Caldara, R., & Schyns, P. G. (2012). Facial expressions of emotion are not culturally universal. Proceedings of the National Academy of Sciences, 109 (19), 7241–7244.

Johnson, E. K., Westrek, E., Nazzi, T., & Cutler, A. (2011). Infant ability to tell voices apart rests on language experience. Developmental Science, 14 (5), 1002–1011.

Juslin, P. N., & Laukka, P. (2003). Communication of emotions in vocal expression and music performance: Different channels, same code? Psychological bulletin, 129 (5), 770.

Kitayama, S., Mesquita, B., & Karasawa, M. (2006). Cultural affordances and emotional experience: socially engaging and disengaging emotions in japan and the united states. Journal of personality and social psychology, 91 (5), 890.

Kotz, S. A., & Paulmann, S. (2007). When emotional prosody and semantics dance cheek to cheek: Erp evidence. Brain research, 1151, 107–118.

Kuhl, P. K., Williams, K. A., Lacerda, F., Stevens, K. N., & Lindblom, B. (1992). Linguistic experience alters phonetic perception in infants by 6 months of age. Science, 255, 606–608.

Meissner, C. A., & Brigham, J. C. (2001). Thirty years of investigating the own-race bias in memory for faces. Psychology, Public Policy and Law, 7 (1), 3–35.

Peirce, J. W. (2007). Psychopy—psychophysics software in python. Journal of neuroscience methods, 162 (1), 8–13.

Pell, M. D., Monetta, L., Paulmann, S., & Kotz, S. A. (2009). Recognizing emotions in a foreign language. Journal of Nonverbal Behavior, 33 (2), 107–120.

Pell, M. D., & Skorup, V. (2008). Implicit processing of emotional prosody in a foreign versus native language. Speech Communication, 50 (6), 519–530.

Perrachione, T. K., Del Tufo, S. N., & Gabrieli, J. D. (2011). Human voice recognition depends on language ability. Science, 333 (6042), 595–595.

Perrachione, T. K., & Wong, P. C. (2007). Learning to recognize speakers of a non-native language: Implications for the functional organization of human auditory cortex. Neuropsychologia, 45 (8), 1899–1910.

Pisoni, D. B. (1993). Long-term memory in speech perception: Some new findings on talker variability, speaking rate and perceptual learning. Speech communication, 13 (1), 109–125.

Ponsot, E., Arias, P., & Aucouturier, J. (2017). Mental representations of smile in the human voice. Journal of the Acoustical Society of America (submitted).

Rachman, L., Liuni, M., Arias, P., Lind, A., Johansson, P., Hall, L., Richardson, D., Watanabe, K., Dubal, S., & Aucouturier, J.-J. (2017). David: An open-source platform for real-time transformation of infra-segmental emotional cues in running speech. Behavior Research Methods, (pp. 1–21).

Russ, J. B., Gur, R. C., & Bilker, W. B. (2008). Validation of affective and neutral sentence content for prosodic testing. Behavior research methods, 40 (4), 935–939.

Sauter, D. A., Eisner, F., Ekman, P., & Scott, S. K. (2010). Cross-cultural recognition of basic emotions through nonverbal emotional vocalizations. Proceedings of the National Academy of Sciences, 107 (6), 2408–2412.

Scherer, K. R., Banse, R., & Wallbott, H. G. (2001). Emotion inferences from vocal expression correlate across languages and cultures. Journal of Cross-cultural psychology, 32 (1), 76–92.

Scherer, K. R., & Oshinsky, J. S. (1977). Cue utilization in emotion attribution from auditory stimuli. Motivation and emotion, 1 (4), 331–346.

Shapiro, P. N., & Penrod, S. (1986). Meta-analysis of facial identification studies. Psychological Bulletin, 100 (2), 139.

Thompson, W. F., & Balkwill, L.-L. (2006). Decoding speech prosody in five languages. Semiotica, 2006 (158), 407–424.

Valentine, T. (1991). A unified account of the effects of distinctiveness, inversion, and race in face recognition. The Quarterly Journal of Experimental Psychology, 43 (2), 161–204.

Van Dillen, L. F., & Derks, B. (2012). Working memory load reduces facilitated processing of threatening faces: An erp study. Emotion, 12 (6), 1340.

Wagner, H. L. (1993). On measuring performance in category judgment studies of nonverbal behavior. Journal of nonverbal behavior, 17 (1), 3–28.

